# Assessment of population differentiation and linkage disequilibrium in *Solanum pimpinellifolium* using genome-wide high-density SNP markers

**DOI:** 10.1101/402420

**Authors:** Ya-Ping Lin, Chu-Yin Liu, Kai-Yi Chen

**Affiliations:** Department of Agronomy, National Taiwan University, Taipei, Taiwan, 10617

**Author notes:** Corresponding author: Kai-Yi Chen Office mailing address: No.1, Sec. 4, Roosevelt Rd., Da’an Dist., Department of Agronomy, National Taiwan University, Taipei City, Taiwan, 10617. **Data available** Short reads of DNA sequences: NCBI SRA BioProject Number PRJNA358110 Supplemental figures: https://doi.org/10.6084/m9.figshare.7010495.v1 Supplemental tables: https://doi.org/10.6084/m9.figshare.7010492.v1.

**Keywords:** *Solanum pimpinellifolium*, population differentiation, linkage disequilibrium pattern, restriction site associated DNA sequencing

## Abstract

To mine new favorable alleles for tomato breeding, we investigated the feasibility of utilizing *Solanum pimpinellifolium* as a diverse panel of genome-wide association study through the restriction site-associated DNA sequencing technique. Previous attempts to conduct genome-wide association study using *S. pimpinellifolium* were impeded by an inability to correct for population stratification and by lack of high-density markers to address the issue of rapid linkage disequilibrium decay. In the current study, a set of 24,330 SNPs was identified using 99 *S. pimpinellifolium* accessions from the Tomato Genetic Resource Center. Approximately 84% *Pst*I site-associated DNA sequencing regions were located in the euchromatic regions, resulting in the tagging of most SNPs on or near genes. Our genotypic data suggested that the optimum number of *S. pimpinellifolium* ancestral subpopulations was three, and accessions were classified into seven groups. In contrast to the SolCAP SNP genotypic data of previous studies, our SNP genotypic data consistently confirmed the population differentiation, achieving a relatively uniform correction of population stratification. Moreover, as expected, rapid linkage disequilibrium decay was observed in *S. pimpinellifolium*, especially in euchromatic regions. Approximately two-thirds of the flanking SNP markers did not display linkage disequilibrium. Our result suggests that higher density of molecular markers and more accessions are required to conduct the genome-wide association study utilizing the *Solanum pimpinellifolium* collection.

## INTRODUCTION

The wild tomato species *Solanum pimpinellifolium* is a native perennial shrub in Ecuador and Peru and is believed to have originated in northern Peru and then diversified into several subpopulations after it migrated to Ecuador and southern Peru (Rick *et al.* 1977; Zuriaga *et al.* 2009; Blanca *et al.* 2012, 2015). The accessions in northern Peru display higher genetic variation and a higher outcrossing rate than those in southern Peru (Rick *et al.* 1977; Caicedo and Schaal 2004). Recent studies suggested that *S. pimpinellifolium* can be divided into at least three subpopulations: one in Peru, one in northern Ecuador and one in the mountains of Ecuador (Zuriaga *et al.* 2009; Blanca *et al.* 2012, 2015). In addition, the major climatic parameters, such as temperature and precipitation, show unidirectional gradient changes from southern Peru towards Ecuador (Zuriaga *et al.* 2009). Because geographic distributions of distinct *S. pimpinellifolium* subpopulations also aligned from south to north, it was proposed that the genetic distances between subpopulations were correlated with climatic differences (Zuriaga *et al.* 2009; Blanca *et al.* 2012, 2015).

*S. pimpinellifolium* is an attractive resource for tomato breeding because it can freely cross with cultivated tomatoes and introduces novel alleles into the limited gene pool of cultivated tomatoes (Tanksley and Mccouch 1997; Spooner *et al.* 2005; Moyle 2008). *S. pimpinellifolium* has been used as a genetic resource for disease resistance and fruit quality traits in tomato breeding (Grandillo *et al.* 2011; Víquez-zamora *et al.* 2014; Capel *et al.* 2015). A core collection of *S. pimpinellifolium* was developed by AVRDC for the purpose of preservation and utilization (Rao *et al.* 2012). This core collection has been used to mine novel alleles of salt tolerance via genome-wide association study (GWAS) (Rao *et al.* 2015).

GWAS utilizes linkage disequilibrium (LD), the non-random association between marker alleles and alleles conferring targeted phenotypes in a given population of germplasm, to map quantitative trait loci (QTLs) (Soto-Cerda and Cloutier 2012). The average ranges of LD decay in different collections of cultivated tomatoes varied from 6 to 13 cM (Sim *et al.* 2012a; Pascual *et al.* 2015; Bauchet *et al.* 2017). It is expected that the range of LD decay is smaller in *S. pimpinellifolium* populations because *S. pimpinellifolium* presents larger genetic variation than cultivated tomatoes (Blanca *et al.* 2012, 2015; Ranc *et al.* 2012; Bauchet *et al.* 2017). Indeed, the SolCAP array with 7,720 SNPs did not achieve full LD coverage across all chromosomal regions for GWAS using *S. pimpinellifolium* accessions (Sim *et al.* 2012a, 2012b; Bauchet *et al.* 2017).

The restriction site-associated DNA sequencing (RADseq) technique might provide an inexpensive solution to address this challenge (Davey and Blaxter 2010). The RADseq technique limits sequencing resources at the vicinity of restriction enzyme cutting sites and therefore provides flexibility of experimental design in terms of the trade-off between cost-effectiveness and marker densities (Chen *et al.* 2014; Bhakta *et al.* 2015).

The objective of the current study was to develop genome-wide high-density SNP markers for a subset of *S. pimpinellifolium* collections from the Tomato Genetic Resource Center (TGRC) through the RADseq approach. The population differentiation and the range of LD decay were then assessed.

## MATERIALS AND METHODS

### Plant materials

All plant materials and their information were obtained from TGRC (Table S1; http://tgrc.ucdavis.edu/). In this study, 12 accessions from Ecuador and 87 accessions from Peru were utilized. According to their mating types, 43 accessions were facultative self-compatible (FSC) and 56 accessions were autogamous self-compatible (ASC). Seeds were propagated by self-pollination for two generations using the method of single-seed descent in a greenhouse. Young leaves collected from plants of these single-seed descendent seeds were used for DNA extraction.

### RAD sequencing

Total genomic DNA was extracted from young leaves using a modified CTAB method (Fulton *et al.* 1995) and purified with a DNeasy Blood & Tissue Kit (QIAGEN, Venlo, Netherland) following the manufacturer’s instructions. *Pst*I-digested DNA libraries were prepared following the protocol of Etter *et al*. (Etter *et al.* 2011). Four RADseq libraries were constructed, and each was sequenced in one lane of an Illumina HiSeq2000 flow cell (100 bp single-end reads) (Illumina Inc., San Diego, CA, USA).

### SNP calling

Reads were analyzed with Stacks version 1.37 (Catchen *et al.* 2013) and with CLC Genomics Workbench software version 6.5.1 (QIAGEN, Venlo, Netherlands) (“CLC Genomics Workbench 6.5.1”). First, the *process_radtags* command in Stacks filtered out low-quality reads with Q scores less than 20. The remaining reads were mapped to the tomato reference genome SL2.50 (Fernandez-Pozo *et al.* 2015) using the “Map Reads to Reference” tool in the CLC Genomics Workbench software. Considering that genetic variation between the tomato reference genome *S. lycopersicum* and *S. pimpinellifolium* is larger than genetic variation within *S. lycopersicum*, mapping parameters were set as 0.5 for the length fraction and 0.9 for the similarity fraction. The reads of the same individual in different lanes were merged together. In the subsequent analyses using Stacks, the *ref_map.pl* command set the parameter –m (minimum read depth to create a stack) as 10, and the *populations* command set the parameter –p (minimum number of populations a locus must be present) as 75. SNPs with a minor allele frequency of less than 0.05 were further excluded, and a set of 24,330 SNP markers was obtained. ITAG2.4 gene model from SGN was used as the reference gene annotation.

### Identification of insertion/deletion (InDel) and simple sequence repeat (SSR) markers

InDels were identified from the Sequence Alignment Map (SAM) files of all *S. pimpinellifolium* accessions with a read depth of no less than two using the “InDels and Structural Variants” tool and the “Compare variants” tool provided in the CLC Genomics Workbench software (“CLC Genomics Workbench 6.5.1”). InDel markers in the form of tandem repeated sequences were classified as SSR markers.

### Population differentiation

To avoid redundant SNP markers used in the subsequent analyses, only one SNP that showed complete LD (*r*^*2*^ = 1) in the same sequencing block around a *Pst*I site was kept whenever more than one SNP existed in the sequencing block. This process resulted in a total of 19,993 SNP markers extracted from the set of 24,330 SNPs noted above. Principle component analysis (PCA) was performed using TASSEL5.0 (Bradbury *et al.* 2007). Population differentiation was investigated via ADMIXTURE (Alexander *et al.* 2009). Calculation of pairwise F_st_ (Weir and Cockerham 1984) and analysis of molecular variance (AMOVA) (Excoffier *et al.* 1992) were conducted in the R packages hierfstat (Goudet and Jombart 2015) and StAMPP (Pembleton *et al.* 2013), respectively.

### Estimate of genetic variation and LD

The genotypes of the 24,330 SNP markers were used to estimate genetic variation and LD in this *S. pimpinellifolium* population. Genetic variation within overall accessions and each of seven groups was assessed based on observed heterozygosity and the within-population gene diversity (expected heterozygosity) using the R package hierfstat (Goudet and Jombart 2015). Pairwise *r*^*2*^ values between SNP markers were calculated to assess overall extent of LD via plink1.9 within a 1-Mb window (Gaunt *et al.* 2007) and fit by non-linear regression (Remington *et al.* 2001). The baseline of the *r*^*2*^ value was set at 0.1 (Bauchet *et al.* 2017). To assess the local LD along each chromosome, we defined the basic unit for local LD as the sequencing region surrounding a *Pst*I site, usually 186 bp long, which has at least one SNP with a minor allele frequency greater than 0.05 in the *S. pimpinellifolium* population. For each pair of consecutive basic units, the average *r*^*2*^ was calculated between two SNPs in different basic units and plotted along the left *Pst*I cutting site based on the physical position. The heterochromatin regions were marked according to the genetic map of EXPIM 2012 and the physical map of the tomato reference genome (Sim *et al.* 2012b).

### Analysis of SolCAP array data of *S. pimpinellifolium*

The SolCAP data of 214 samples of *S. pimpinellifolium* were downloaded from previous studies (Blanca *et al.* 2012, 2015; Sim *et al.* 2012a). A set of 4,326 bi-allelic SNPs was first extracted and filtered with the criteria that minor allele frequency is greater than 0.05 and the proportion of missing genotypes is less than 25%. These SNP-filtering criteria are the same as the criteria applied to the SNP dataset generated in this study. This procedure resulted in 2,817 SNPs. Subsequently, population differentiation was investigated by ADMIXTURE (Alexander *et al.* 2009). Because some accessions appeared in different SolCAP genotyping studies and their genotypes were not completely matched, different suffixes—“_2012S,” “_2012B,” and “_2015B,”—were added to the accession name to indicate their origins from references Sim *et al.* 2012a, Blanca *et al.* 2012, and Blanca *et al.* 2015, respectively. In addition, the percentage of identical SNP genotypes between accessions with the same name was calculated based on the 4,326 SNP genotypes and excluding missing values.

### Data avalibility

All the sequences of RADseq are available at the NCBI SRA database, and the BioProject Number is PRJNA358110. Supplemental files available at FigShare: https://doi.org/10.6084/m9.figshare.7010495.v1 for supplemental figures; https://doi.org/10.6084/m9.figshare.7010492.v1 for supplemental tables.

## RESULTS

### Identification of 24,330 SNPs from *Pst* I-digested DNA libraries

A total of 655,973,270 short DNA reads were obtained from four lanes of the Illumina HiSeq2000 flow cell and were divided into 99 parts according to barcode sequences. Each part was derived from the DNA of a *S. pimpinellifolium* accession and contained at least 3.7 million DNA reads, except for LA2647 (Table S1). To ensure the accuracy of SNP calling and genotype calling, two criteria were set: one was that the read depth aligning to the reference sequences was equal to or greater than 10, and the other was that at least 75% of the accessions showed genotypes associated with a defined SNP marker.

Among the 82,814 *Pst*I sites in the tomato reference sequence SL2.50, only 23,988 *Pst*I sites were around the sequenced DNA reads (Table S2). The sequenced regions included 0.54% of the SL2.50 reference sequences and 12,790 annotated genes (Table 1). Interestingly, approximately 84% of the sequenced *Pst*I sites were located in the euchromatic regions (Table S2). Nevertheless, no significant difference was observed in the proportion of sequencing regions for SNP discovery between the euchromatic regions (68.85%) and the heterochromatic regions (60.59%) (Table S2). A total of 67,804 SNPs were identified in the sequenced regions of 99 *S. pimpinellifolium* accessions, and 24,330 of them had a minor allele frequency greater than 0.05.

In the genotypic dataset of 24,330 SNP markers (Table S3), the missing proportion of each accession ranged from 0.72% to 15.92%, except for LA2647, for which the value was 65.68% due to a low number of sequencing reads (Table S1). Regarding the features of these 24,330 SNPs, 16,365 SNPs were located in 7,383 annotated genes (Table 1) and the remaining SNPs were located in the intergenic regions. In addition, 3,068 InDels (Table S4) and 107 SSR markers (Table S5) were obtained. In the subsequent analyses, only SNP markers were utilized, and the genotypic data of the LA0411 accession was dropped because the observed heterozygosity of LA0411 was inconsistent with its mating type (Table S1).

### Genetic differentiation of *S. pimpinellifolium* corresponded to the geographic area

The collection of 98 *S. pimpinellifolium* accessions in this study was divided into seven groups corresponding to three ancestral populations using the ADMIXTURE software (Figure 1A and Figure S1). The seven groups included three groups with pure ancestry, three groups with an admixture of two different ancestries, and one group with an admixture of three ancestries. As expected, accessions in each group were clustered together in the PCA plot, in which principal component 1 (PC1), PC2, PC3, PC4 and PC5 explained 16.04%, 8.00%, 3.94%, 3.12% and 2.54% of the variation, respectively (Figure 1B). Interestingly, most accessions in the same group were in the same vicinity in terms of their collection sites (Figure 1C). In addition, different ancestral groups were spread in somewhat distinct geographic areas along the coastline from Ecuador to southern Peru (Figure 1C). The geographic distribution of these groups appeared in the following order from north to south: the pure red ancestral group, the admixture group with red-blue ancestries, the pure blue ancestral group, the admixture group with blue-green ancestries, and the pure green ancestral group (Figure 1C). This geographic distribution showed a trend in which the admixture groups were located between their corresponding pure ancestral groups.

**Figure 1.**
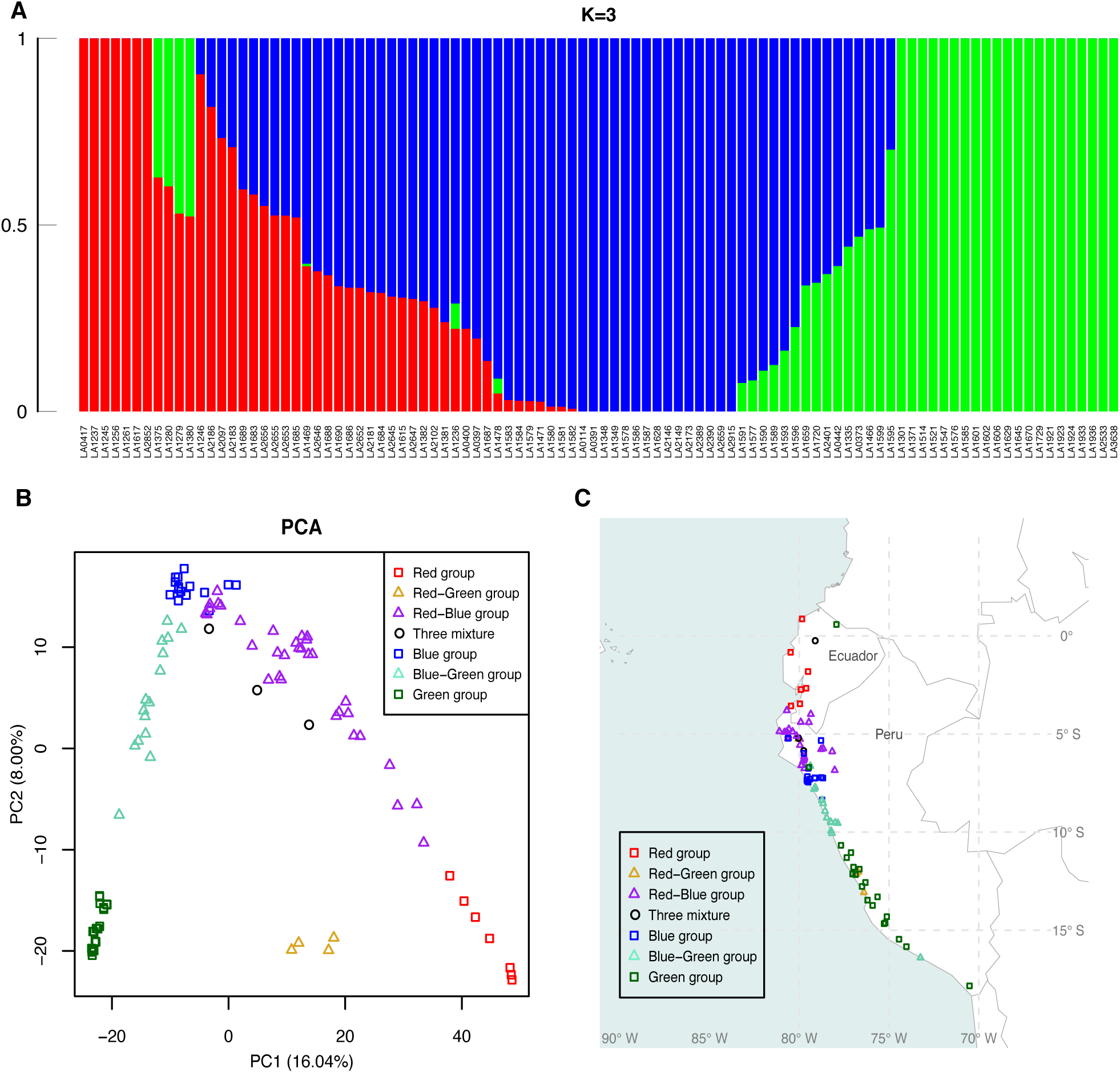
Ancestry and geographic distribution of 98 *Solanum pimpinellifolium* accessions from the Tomato Genetics Resource Center. A) Model-based ancestry for each accession. B) Principle component analysis of the *S. pimpinellifolium* population. C) Geographical distribution of the 98 *S. pimpinellifolium* accessions. Symbol and color codes are as follows: square symbols with red, blue and green colors were used to indicate three pure ancestry groups corresponding to the same colors in the ancestry plot; triangle symbols with goldenrod, purple and aquamarine colors were used to present the three admixture groups with red-green, red-blue and blue-green mixing ancestries, respectively; black circle symbols were used for the group with admixture of three ancestries.

To compare genetic variation within pure ancestral groups or within admixture groups, the within-population gene diversity of each group was calculated. The blue group and the red-blue group showed the highest genetic variation among the pure ancestral groups and the admixture groups, respectively (Table 2). Both groups were in northern Peru, which indicated that northern Peru is the origin of *S. pimpinellifolium*. Pairwise F_st_ confirmed the population differentiation (Table S6), and AMOVA revealed that the variation between groups was 41.96% (p-value < 0.001).

The differentiation of two mating types, FSC and ASC, was expected because non-random mating would disrupt the Hardy-Weinberg equilibrium, leading to population structure (Weir and Cockerham 1984; Holsinger and Weir 2009). In this collection, most FSC accessions were clustered in northern Peru, while ASC accessions were scattered in Ecuador and central and southern Peru, along the western side of the Andes Mountains to the coast (Figure S2). The pairwise F_st_ (0.0029) of FSC and ASC was significant (p-value < 0.001). However, PCA presented unclear clusters between FSC and ASC (Figure S3). In addition, the variation between FSC and ASC was only 5.91% despite the significance of AMOVA (p-value < 0.001).

### Rapid LD decay

Overall LD decay was estimated for the mapping resolution in GWAS. In this population, the non-linear regression curve dropped very quickly (Figure S4). Following the non-linear regression curve, the overall LD decay was within 18 Kb when the baseline of the *r*^*2*^ value was set at 0.1 (Table 3 and Figure 2A). The fastest LD decay was within 10 Kb on chromosome 9 while the slowest decay was within 30 Kb on chromosome 4 (Table 3 and Figure S5).

**Figure 2.**
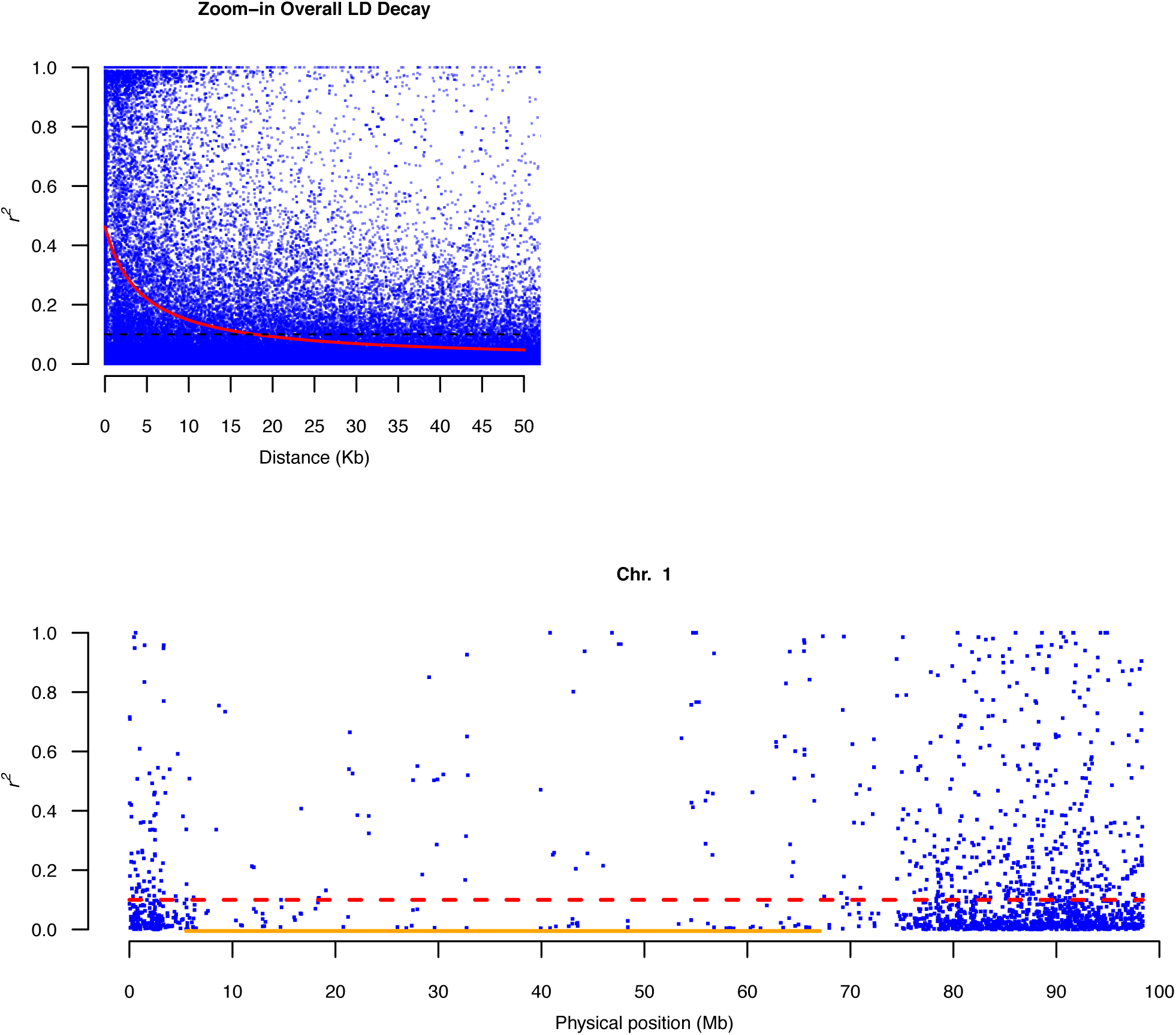
Visualization for LD. A) The 50 Kb interval of overall LD decay. The red curve indicates non-linear regression, and the black dotted line indicates the baseline of *r*^*2*^ at 0.1. B) The local LD of chromosome 1. The red dotted line indicates the baseline of *r*^*2*^, and the orange line indicated the heterochromatic region.

### Heterogeneity of genetic recombination within each chromosome

LD decay has often been estimated for each chromosome (Sim *et al.* 2012a; Bauchet *et al.* 2017). However, the LD decay per chromosome was insufficient to capture the local variations of historically accumulated recombination events because the tomato genome comprises more than 75% heterochromatin, which usually suppresses recombination events (Sim *et al.* 2012a). The local LD profile of individual chromosomes was assessed based on the average *r*^*2*^ value of flanking sequencing units that contained at least one SNP marker. Two major trends were observed (Figure 2B and Figure S6). Marker density in the heterochromatic regions was lower than that in the euchromatic regions, and approximately two-thirds of the *r*^*2*^ values were less than 0.1 (Table 3). The latter observation indicated that these flanking SNP markers were not in a state of linkage disequilibrium.

## DISCUSSION

### A similar distribution between genes and SNPs was identified in the vicinity of *Pst*I cutting site throughout the genome

The observation that 67.26% (16,365 to 24,330) of the SNPs were located in the annotated gene regions (Table 1) implied a correlation between the distribution of the identified SNPs in the current study and the distribution of the annotated genes. Additional observations in the current study indicated a preference for genomic DNA digestion by the *Pst*I restriction enzyme in the euchromatic regions: only 28.97% (23,988 to 82,814) of *Pst*I sites were found in the deep sequencing regions, and 83.55% (20,043 to 23,988) of the deep sequencing regions were located in the euchromatic region (Table S2). It is worth noting that the current RADseq protocol did produce low coverage of sequencing reads in certain *Pst*I sites (with a read depth less than 10), and these *Pst*I sites were filtered by the criteria of SNP and genotype calling; therefore, the deep sequencing regions indicated that their read depths were no less than 10. Incidentally, because SNPs can be identified only in the sequenced regions, it is a reasonable deduction that most SNPs found in the current study are located in the euchromatic regions. Plotting the annotated genes, the expected *Pst*I sites, the *Pst*I sites in the deep sequencing regions, and the 24,330 SNPs identified in the current study (Figure 3A, 3B, 3C, and 3D, respectively), shows clearly that the annotated tomato genes, the *Pst*I sites in the deep sequencing regions, and identified SNPs are mainly located in the euchromatic regions.

**Figure 3.**
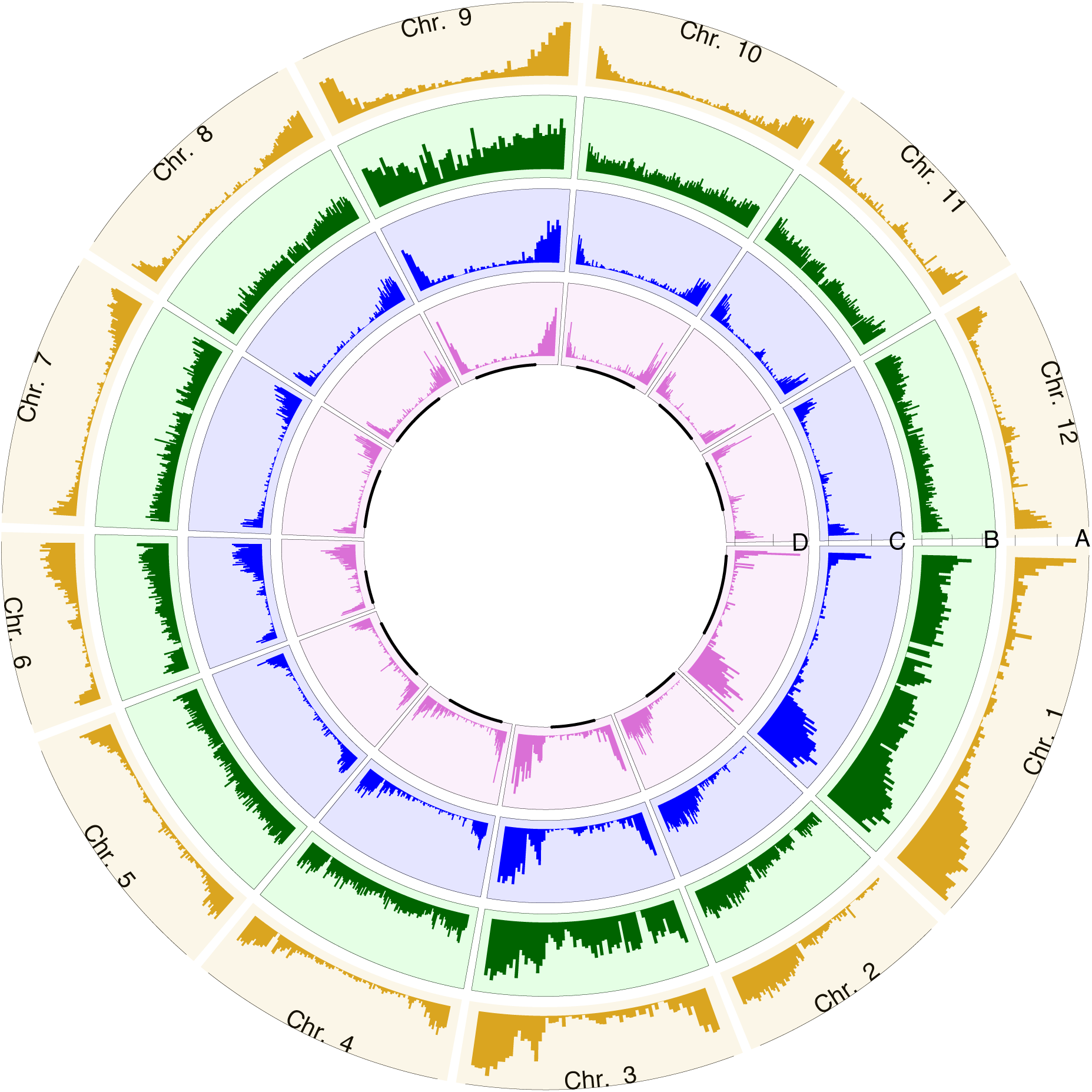
The distributions of ITAG2.4 gene model, *Pst*I cutting sites and SNPs throughout the genome. Each section indicates one chromosome, with labeling on the circumference. Circles A, B, C, and D indicate the distribution of ITAG2.4 genes, expected *Pst*I cutting sites, *Pst*I cutting sites in the deep sequencing regions and RADseq SNPs, respectively. The black lines in the inner D layer indicate the heterochromatic regions.

*Pst*I is a methylation-sensitive restriction enzyme and recognizes the sequences “CTGCAG” (Dobritsa and Dobritsa 1980). The study of the genome-wide methylation pattern in tomato leaves and immature fruits revealed that the gene-rich euchromatic regions at the distal ends of chromosomes were characterized as the regions with low levels of cytosine methylation at the “CG”, “CTG”, and “CAG” sequences, and the pericentromeric heterochromatin regions were the regions with high levels of cytosine methylation (Zhong *et al.* 2013). Because the young tomato leaves were used as the DNA source to construct the RADseq libraries, it is reasonable to infer that the *Pst*I-digested RADseq-targeted chromosomal regions were concentrated in the gene-rich euchromatic regions. Therefore, one can emphasize the sequencing resources on euchromatic regions via *Pst*I RADseq when preparing candidate gene research for tomatoes.

### The discrepancy in inferences of population differentiation of *S. pimpinellifolium*

The estimation of the best subpopulation number (K) is very important in GWAS because population structure is integrated as a correction to eliminate the inflated significance due to confounding effects (Korte and Farlow 2013). If the best K of a certain population could not be confirmed, the results of GWAS would be unreliable. However, several previous studies did not achieve the same best K of *S. pimpinellifolium*: 10 SSR markers for 248 individuals obtained an unclear K (Zuriaga *et al.* 2009); 48 SSR markers for 190 individuals revealed a best K = 2 with admixtures following a K = 5 (Rao *et al.* 2012); finally, the SolCAP array for two collections of 63 and 112 individuals obtained the same best K = 3 with admixtures (Blanca *et al.* 2012, 2015). Our study obtained the best K = 3, but our ancestral and admixture groups were different from the latter studies of the SolCAP array (Blanca *et al.* 2012, 2015). These previous studies suggested that the *S. pimpinellifolium* population was differentiated into three ancestral groups: one in the northern Ecuador; another in the mountainous area from southern Ecuador extending to northern Peru, and the third in the low-altitude areas of Peru, along with certain admixtures (Blanca *et al.* 2012, 2015). In contrast, our study showed that the *S. pimpinellifolium* accessions were clustered into three pure ancestral groups, with one in Ecuador (the red group), another in northern Peru (the blue group), and the third in southern Peru (the green group), as well as three clearly identified admixture groups (Figure 1C).

To investigate the potential reasons for the inconsistent conclusions between the current study and the previous studies based on SNP markers, genotypic data of the *S. pimpinellifolium* accessions made from the SolCAP array in three previous studies were obtained from internet (Blanca *et al.* 2012, 2015; Sim *et al.* 2012a) and a meta-analysis was conducted using our workflow (please see details in the “Materials and Methods” section) (Table S7). A total of 214 samples representing 126 accessions were divided into 11 groups via ADMIXTURE using filtered genotypes of 2,817 SNP markers.

However, the results of this meta-analysis pose two problems. First, the cross-validation error did not confirm that K = 11 was the optimal grouping method (Figure S7 and Figure S8). This condition can be explained as a low population structure in a population with high genetic diversity, which may have resulted from frequent gene flow (Gevaert *et al.* 2013).

Overrepresentation of common SNPs within *S. pimpinellifolium* accessions on the SolCAP genotyping array could be the other reason given that the SolCAP array was originally created to explore the genetic variation within cultivated tomatoes and to map genes (Hamilton *et al.* 2012; Sim *et al.* 2012a, 2012b).

The second problem was that certain samples belonging to the same accession were not clustered in the same group. For example, two samples of the BGV007104 accession, which shared 93.23% genotypic identity in this SNP set (Table S8) and were labeled as BGV007104_2012B and BGV007104_2015B, were assigned to different groups (Figure S8). The wrong grouping for the samples of the same accession may result from an insufficient number of SNP markers that were unable to capture similarity within the same accession when the sample size increases. In an empirical study of *Arabidopsis halleri*, a few thousand SNP markers were required to estimate the genetic diversity among populations with different genetic variation (Fischer *et al.* 2017). The latter problem prevented meaningful comparisons between the inference of population differentiation in the current study and that of the meta-analysis. Regardless of the inconsistent best K among these studies, the results of pairwise F_st_ and AMOVA statistically supported the subpopulations in this study, suggesting that this set of high-density SNPs could stably estimate the best K.

### A group of individuals would be a better representative of an accession

An accession should be represented by a group of samples rather than only a single individual since an accession in its natural habitat is composed of a group of individuals, especially when gathering accessions with high diversity. However, under circumstances with limited resources, we instead prepared a collection representing a population rather than only several accessions because our final goal was to apply *S. pimpinellifolium* in GWAS. In addition, the mating system and the propagation method of *S. pimpinellifolium* made the variations between accessions greater than that within accessions. Therefore, our only option was to involve as much diversity as possible to enhance the efficiency of GWAS.

### High genetic variation leads to rapid LD decay

The observed and expected heterozygosity of this population were 0.0761 and 0.2786, respectively, slightly higher than in previous research (Blanca *et al.* 2012, 2015). Since *S. pimpinellifolium* was detected with up to a 40% outcrossing rate (Rick *et al.* 1977) and demonstrated high genetic variation, it is expected to cause rapid LD decay. In this study, LD decay was within 18 Kb throughout the genome, which was much shorter than in cultivated tomatoes (Sim *et al.* 2012a; Bauchet *et al.* 2017). However, such high genetic variation requires much more markers to enable the comprehensive detection in GWAS. The 900-Mb tomato genome requires at least 50,000 markers to cover the entire genome evenly. Therefore, acquiring more SNPs using different methods is essential to conduct a GWAS in the *S. pimpinellifolium* population. One possible approach is to increase the sample size evenly for each subpopulation (Brachi *et al.* 2011). Since approximately 64% of alleles were rare in this population, the augmentation of the subpopulation size may adjust rare alleles to common alleles, potentially increasing the SNPs without extending coverage. Another possible strategy is exome sequencing, a selective genome sequencing technology that selects desired sequencing regions by the hybridization of designed probes (Kaur and Gaikwad 2017). Based on tomato genome sequence information, such as the gene model or EST database, one could design different sets of probes to limit sequencing regions (Ruggieri *et al.* 2017). Given the approximately 110 Mb total gene length in the ITAG2.4 gene model, the potential coverage could reach 12% and all target the gene region. This exome sequencing strategy may be able to increase SNPs without increasing the population size.

### A reproductive strategy would reduce the genetic diversity of *S. pimpinellifolium*

These accessions were propagated using single-seed descent for two generations. Therefore, the heterozygosity would be reduced compared to the original specimens, especially for FSC accessions. Here, we revealed that an ASC accession, LA0411, presented 40.25% heterozygosity, which highlighted the contradiction of self-fertilization consequence. Lacking the same accession as a reference in previous studies and considering the 0 to 22% heterozygosity of other accessions in the original published research (Rick *et al.* 1977), we could remove only LA0411 from our analyses based on the fact that its heterozygosity was too high for an ASC accession.

## ACKNOWLEDGMENTS

We thank the C.M. Rick Tomato Genetics Resource Center for generously providing plant materials to conduct this experiment and Dr. Tze-Tze Liu in the Genome Research Center at the National Yang-Ming University, Taiwan, for providing help and advice of the massive parallel DNA sequencing task. This work was supported by grants from the Ministry of Science and Technology, Taiwan R.O.C. (Project Nos. NSC 96-2313-B-002-034-MY3 and NSC 101-2313-B-002-006-MY3) to KYC and from National Taiwan University (Project Nos. 103R7858 and 104R7858) to KYC. This work represents the partial fulfillment of requirements for YPL’s Ph.D. degree.

